# Rainbow trout neomales broodstocks are able to eat and use a high carbohydrate diet during a complete reproductive cycle

**DOI:** 10.1101/2023.02.20.529192

**Authors:** Favalier Nathan, Surget Anne, Véron Vincent, Maunas Patrick, Turonnet Nicolas, Terrier Frederic, Panserat Stéphane, Marandel Lucie

**Author notes:** **Corresponding author:** Dr L Marandel, INRAE, Université de Pau et des Pays de l’Adour, e2s, St-Pee-sur-Nivelle, France.

## Abstract

The main challenge of the aquaculture relies on the shift from fishmeal to more sustainable ingredients. Interest on carbohydrates has growing rapidly as they are considered as promising resources, providing energy and preserving proteins for growth. However, rainbow trout is considered as a poor user of carbohydrates displaying postprandial hyperglycaemia and impaired growth performances when fed with diets containing more than 20% of carbohydrates. Nonetheless, recent evidence points that broodstocks could be better user of high carbohydrate diet compared to juveniles. However, no study investigates how sex-reversed females – neomales – eat, grow and reproduce under a high carbohydrate diet. Our objective was thus to assess growth and reproductive performances of neomales fed with a high carbohydrate diet during an entire reproductive cycle. Our results demonstrate that neomales display specific metabolic and physiological changes when fed with a high carbohydrate diet compared to both females and males broodstocks as well as compared to juveniles. Altogether, our data demonstrate the critical relevance to formulate specific diets in accordance with specificities of each type of broodstocks (*i*.*e*. females, males and neomales).

## 1 Introduction

Aquaculture is growing rapidly, and its production become integral in global food resources, providing a large part of the world’s seafood supply. However, the production of fish meal (FM) and fish oil (FO), that are traditional ingredients in aquafeed relying on world capture fisheries, remains limited and hinders a sustainable development of aquaculture. Thus, one of the main challenges for the aquaculture industry rely on the shift from fishmeal (FM) to other sources of proteins and tremendous effort have been made to achieve this [1]. In this regard, interest on carbohydrates that can be considered as promising resources has grown rapidly both in the scientific and fish farmer communities. Digestible carbohydrates represent a non-negligible source of energy for fish and preserve proteins for growth [2] as well as provide ecological benefits by reducing nitrogen excretion. In addition, plant-derived carbohydrates can be considered in European region as easy-to-produce and inexpensive raw material [3]. The relevance of the use of plant-derived carbohydrates is all the more important when feeding broodstocks fish, as they require a large quantity of expensive feed to be maintained and reproduced. However, when the substitution of FM by digestible carbohydrates exceeds 20% in the aquafeed formula of rainbow trout, juveniles display a decrease of growth and have a persistent postprandial hyperglycaemia. Several hypotheses have been stated to explain this phenotype. First, on one hand it could be linked to the non-inhibition of the production of endogenous glucose by the liver through gluconeogenesis [4]. On the other hand, it could be linked to the poor induction of the *de novo* lipogenesis responsible of the conversion of glucose into free fatty acids which has been described in juveniles whereas the use anti-diabetic drugs (metformin) lead to a better postprandial glycaemic control through the activation of this pathway [5]. Moreover, the poor use of carbohydrates could be also linked to the weak potential of glucose to induce insulin secretion through for instance poor muscles’ capacity to phosphorylate glucose associated with low numbers of glucose transporters and/or insulin receptors in these tissues [6–9].

Nonetheless, most of the studies regarding the glucose metabolism has been performed on juveniles from the first feeding to commercial size but less interest has been given to broodstock fish. Callet *et al*., 2020 reported that both males and females broodstocks were able to eat and use a high-carbohydrate diet (35%) during all the reproductive cycle (9 months) for females and the early part for males (4 months) as reflected by a conserved growth and good reproductive performances. Both females and males were never hyperglycaemic when fed enriched-carbohydrate diet, demonstrated by specific regulation of both the glucose and lipid metabolisms, compared to what is observed in juveniles. However, only the long-term consequences of feeding females and males with a high carbohydrate diet were evaluated. Nonetheless, a third sex is used in rainbow trout aquaculture, called neomales. Neomales are sex-reversed females used to produce all-female population. They can be obtained through the addition of α17-methyltestosterone in the feed during three months after the first feeding. This broodstock undergo one complete reproductive cycle and generally are sacrificed to collect the gametes during the reproductive event. Thus, neomales rainbow trout are mainly studied for their sperm quality and quantity, fertilisation performances or the gonadal structure throughout the reproductive cycle [10–12]. However, as far as we know no studies focussed on their nutrition and potential effects of using specific formulated feed on their growth and reproductive performances. Recently, we demonstrated that a short period of glucose intake (two day – 5 meals) did not induce specific metabolic in mature female and male and neomale [13]. However, compared to both mature females and males, neomales displayed more pronounced modifications of their metabolism in response to nutritional status and HC diet (especially a huge increase in Gck mRNA levels)[13]. Although this could be attributed to their physiological status and weight close to juveniles during the experiment, it cannot be excluded that neomales are better user of high carbohydrate diet compared to juveniles.

In this context, the aim of the present study is to assess long-term consequences of the use of a high-carbohydrate diet on the growth and reproductive performances as well as on the hepatic and gonadic metabolism of neomales.

## 2 Materials and methods

### 2.1 Ethics approval

Experimentation was conducted according to the guiding principles for the use and care of laboratory animals and in compliance with French and European regulations on animal welfare (Décret 2001-464, 20 May 2001, and Directive 2010/63/EU, respectively).

### 2.2 Fish diets

Neomales rainbow trout were obtained through the addition of 17 -methyltestosterone (3mg.g-1) in the feed during three months after the first feeding at the experimental facility of PEIMA (Sizun, France) which result in 100% of masculinisation [14]. Two experimental diets for rainbow trout, i.e., NC (high-protein and low-carbohydrate diet) and HC (medium-protein and high-carbohydrate diet), were prepared in our own facilities (INRAE, Donzacq, Landes, France; https://doi.org/10.15454/GPYD-AM38) as extruded pellets. Gelatinised starch was included as the carbohydrate source and protein was originated from fish meal and dietary lipids from fish oil and fish meal. The amount of carbohydrates (∼33%) was balanced by the amount of protein. Chemical composition of diets was analysed through several steps: Kjeldahl method after acid digestion was used to determine crude proteins content, gross energy was measured in an adiabatic bomb calorimeter (IKA, Heitersheim Gribeimer, Germany), dry matter was assessed after drying to constant mass at 105°C, crude lipids by petroleum ether extraction (Soxtherm) and ash was estimated through incineration in a muffle furnace for 8h at 550°C.

### 2.3 Experimental design

Neomales used in this experiment were autumnal strain that reproduce during winter in France (November-December) and were obtained through sex-reversion of females by the addition of 3mg.g-1 of α17-methyltestosterone in the diet during three month after the first meal. Two-year-old neomales were fed with either the NC or the HC diet at the INRAE Lées-Athas experimental fish farm at 8°C (figure 1; https://doi.org/10.15454/GPYD-AM38). Due to the large number of fish and the duration of the experiment, we distributed the fish in one tank per condition (NC or HC). The feeding trial was conducted from the resumption of feeding after the reproduction period (December 2019) to the next reproduction period (November 2020). Fish were sampled in February (Figure 1 -Sampling 1), June (Sampling 2), August (Sampling 3) and November 2020 (Sampling 4). During each sampling event, 9 neomales per diet were anaesthetized in a benzocaine bath at 30 mg.L-1 and then killed in a benzocaine bath at 60 mg.L-1. For the first samplings in February, June and August fish were sampled 6h after the last meal to complete the digestion process and obtain postprandial effect of the last meal while in November fish were fasted prior to the reproductive event. Blood was each time drawn from the caudal vein and centrifuged at 3000xg for 15min. Plasma was then recovered and stored at -20°C for future plasmatic analysis. Livers and gonads were dissected into three parts, two parts were snap frozen in liquid nitrogen for RNA extraction and enzymatic activities while the last part was kept at -20°C for biochemical analysis. Gonads were not present in every neomales during all our samplings: in February no neomales had gonads, in June 7 NC neomales and 5 HC neomales had gonads and in August and November 5 of both NC and HC neomales had gonads. Viscera (stomach, intestine and digestive tract) and the remaining carcass (without the liver, gonads and viscera) were also kept at -20°C for biochemical analysis. During the spawning period in November 2020, spawns from 6 females fed with the NC diet were cross-fertilized with milts from 6 neomales from each experimental condition (NC and HC diets). Each of the 70 crossings were closely monitored for survival at eyed stage and at hatching, and for the presence of malformations.

**Figure 1:**
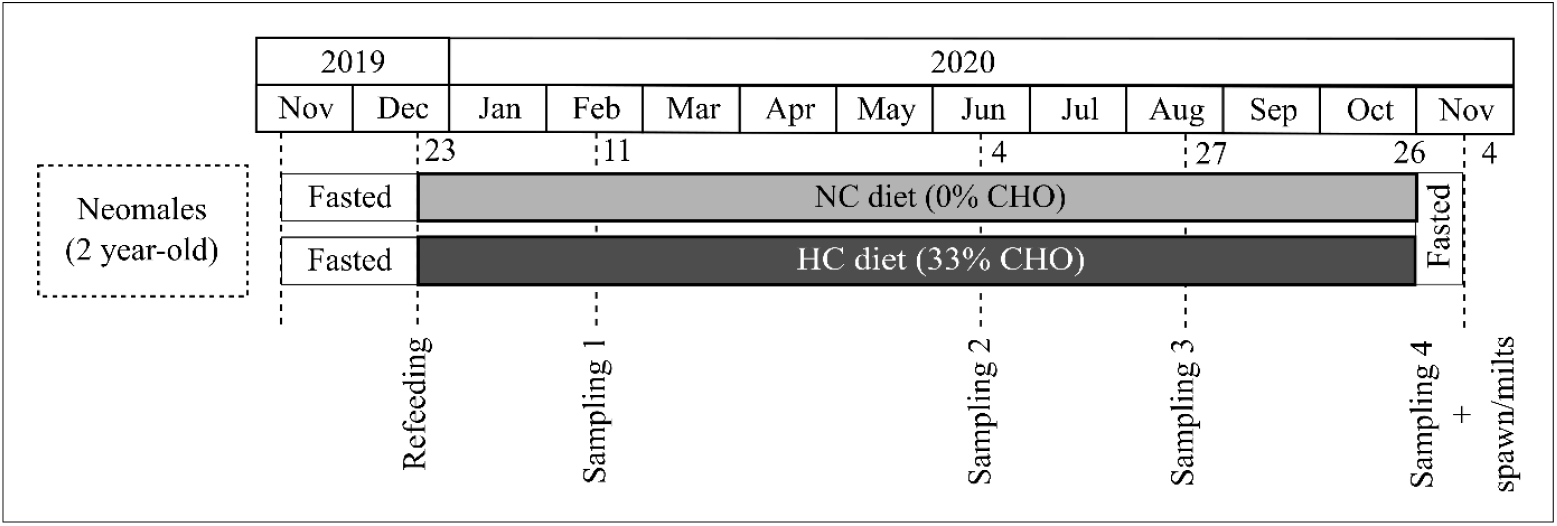
Experimental design. Neomales were fed with a high-carbohydrate diet (HC – 33%) and a non-carbohydrate diet (NC) from December 2019 to November 2020 and the spawning period of females of the same strain.

### 2.4 Biochemical composition of gonads, livers, viscera and carcasses

Glycogen contents were analysed on lyophilized organs. Glycogen content was determined by a hydrolysis technique previously described by Good et al. (1933) [15]. Each sample was homogenised in 1 mol.L-1 HCl (VWR, United States). Free glucose content was measured at this stage. After 10min centrifugation at 10000xg free glucose was measured using the Amplite Fluorimetric Glucose Quantification Kit (AAT Bioquest, Inc., United States) according to the manufacturer’s instructions. The remaining ground tissue was boiled at 100°C for 2.5h and then neutralized by 5 mol.L-1 KOH (vWR, United States). Total glucose (free glucose + glucose obtained from glycogen hydrolysis) was measured with the same kit. Glycogen content was evaluated by subtracting free glucose levels. Total lipid content was determined gravimetrically after extraction by dichloromethane/methanol (2:1, v/v), containing 0.01% of butylated hydroxytoluene (BHT) as antioxidant, according to Folch et al (1957). Total protein was measured using the Kjeldahl method.

### 2.5 Plasma metabolites

Plasma glucose and triglycerides concentrations were measured with Glucose RTU (BioMerieux, Marcy l’Etoile, France) and PAP 150 (BioMerieux) kits, respectively, according to the recommendations of each manufacturer.

### 2.6 RNA extraction and cDNA synthesis

Nine neomales per diet and per sampling (NC and HC) were used for RNA extractions. Liver and gonads (when present) samples were homogenised using Precellys®24 (Bertin Technologies, Montigny-le-Bretonneux, France) in Trizol reagent (Invitrogen, Carlsbad, CA, USA) with 2.8mm ceramic beads for 2×20s, separated by 15s of break, at 5500 rpm. Total RNA was extracted according to the instructions of the manufacturer of Trizol reagent. Total RNA obtained (2μg) was then reverse transcribed to cDNA in duplicate using the SuperScript III RNase H-Reverse Transcriptase kit (Invitrogen) with random primers (Promega, Charbonnières-les-bains, France).

### 2.7 Quantitative real-time PCR in livers and testis

Quantitative real-time PCR (qPCR) was used to measure mRNA levels of genes pertinent to (1) glucose transport, with *glut2a* and *glut2b* both coding for the glucose transporter 2, *glut1aa, glut1ab, glut1bb* and *glut1ba* coding for the glucose transporter 1 and *glut4a* and *glut4b* coding for the glucose transporter 4 a; (2) glycolysis with *gcka* and *gckb* coding for the glucokinase, *pfkla* and *pfklb* coding for the phosphofructokinase and pkl coding for the pyruvate kinase; (3) gluconeogenesis with *g6pca, g6pcb1a, g6pcb1b, g6pcb2a* and *g6pcb2b* coding for the glucose-6-phosphatase, *fbp1a, fbp1b1* and *fbp1b2* coding for the fructose-1,6-biphosphatase and *pck1, pck2a* and *pck2b* coding for the phosphoenolpyruvate kinase; (4) pentose phosphate pathways and *de novo* lipogenesis with *g6pd* coding for the glucose-6-phosphate dehydrogenase, *srebp1* coding for the sterol regulatory element-binding protein 1, acly coding for the ATP-citrate lyase, *aca alpha a* and *aca alpha b* coding for the isoform of the acetyl-coenzyme A carboxylase. Primers sequences used in real-time RT-PCR assays for gluconeogenic, glycolytic and lipogenic genes analysis were those previously described by Marandel et al. [4] and Liu et al. [16]. qPCR was also used to measure mRNA levels specifically in gonads pertinent to (1) steroidogenesis with *fshr, lhcgr* and *star* coding respectively for the follicle stimulating hormone receptor, luteinizing hormone/choriogonadotropin receptor and steroidogenic acute regulatory protein and (2) spermatogenic maturation markers with *nanos2* coding for the nanos C2HC-type zinc finger 2, *dazl* coding for the deleted in azoospermia like and *plzfb* coding for the Zinc finger and BTB domain containing 16b. Primers sequences used in qPCR assays for steroidogenesis and spermatogenic maturation were described respectively by Sambroni et al. (2013) [17] and Bellaiche et al. (2014) [18].

Real-time RT-PCR were performed with the Roche Lightcycler 480 system (Roche Diagnostics, Neuilly-Sur-Seine, France). The mix of the reaction was 6 μL per sample : 2 μL of diluted cDNA (1:76), 0.24 μL of each primer (100 mmol.L-1), 3 μL of Light Cycler 480 SYBR Green 1 Master mix and 0.52 μL of DNase / RNase-free water (5 Prime Gmbh, Hamburg, Germany). Initiation of the real-time RT-PCR protocol was at 95°C for 10min for the denaturation of the cDNA and hot-start Taq-polymerase activation, followed by 45 cycles of a two-step amplification program (15s at 95°C; 10s at 60°C). Melting curves were verified systematically (temperature gradient 1.1°C/15s from 65 to 97°C) at the end of the last amplification cycle to confirm the specificity of the reaction of amplification. Each assay comprised replicates (duplicate for both reverse transcription and PCR amplification) and negative controls (reverse transcriptase-and cDNA template-free samples). Data were then normalised to the 18S (forward 5′-CGGAGGTTCGAAGACGATCA-3′, reverse 5′-TCGCTAGTTGGCATCGTTTAT-3′) transcript abundance in samples of liver. A geometric mean of the following genes as previously described by [19] : 18S, ef1α (forward 5′TCCTCTTGGTCGTTTCGCTG-3′, reverse-5′-ACCCGAGGGACATCCTGTG-3′), β-actin (forward 5′-GATGGGCCAGAAAGACAGCTA-3′; reverse 5′-TCGTCCCAGTTGGTGACGAT-3′) and RS16 (forward 5′-CACAACCATGGGCAAGAAGA-3′, reverse 5′-TCAGGGCAGGGTCTCTGAAG-3′) was used for gonads samples.

### 2.8 Enzymatic activities in the liver

Enzymatic activities were assessed with frozen samples of liver. Samples were ground in an ice-cold buffer (50 mmol.L-1 TRIS, 5 mmol.L-1 EDTA, 2 mmol.L-1 DTT, protease inhibitor cocktail (Sigma, St. Louis, MO, USA)). For the glucose-6-phosphate dehydrogenase (G6pd) and the fatty acid synthase (Fasn) samples were centrifuged for 20 min at 24000g at 4°C and supernatants were kept for the enzymatic assay. To measure phosphofructokinase (Pfk) and pyruvate kinase (Pk) activities samples were centrifuged again for 20 min at 20000g at 4°C. For the glucose-6-phosphatase (G6pc), a sonic disruption (Bioruptor, 3 cycles, 30 s On / 30 s Off), followed by a second centrifugation (10 min at 900xg at 4°C) and supernatants were kept for the enzymatic assay. Enzymatic activities were measured following the variation if the absorbance of nicotinamide adenine dinucleotide phosphate at 340 nm, in a Power Wave X (Biotek Instrument) reader. Reactions were started by the addition of specific substrates. Water was used as a blank for each sample. The enzymes assayed were : high km hexokinase (Gck) following the protocol of [20], fructose-6-bisphosphatase (Fbp) as described in [21], phosphofructokinase (Pfk) according to [22], fatty acid synthase (Fasn) according to the protocol of [23], G6pd and G6Pase respectively according to [24] and [25].

### 2.9 Statistical analysis

R software (v3.6.1) was used to perform all statistical analysis. Results are presented as mean ± SEM (Standard error mean). Statistical analyses were carried out on the different parameters recorded to investigate potential differences among groups of NC and HC neomales. When the normality of distributions and homogeneity of variances were confirmed, two-way ANOVA were performed on the different variables obtained. If conditions of application of a two-way ANOVA were not verified, Mann-Whitney test were performed. When significant interaction (Date and Diet) was demonstrated, means were compared using a Tukey’s post hoc analysis. Statistical difference was considered significant when p-value was <0.05.

## 3 Results

### 3.1 Chemical body composition

Body weight as well as body composition of neomales were assessed after each sampling (table 2). Mean body weight of neomales were globally significantly different among month of sampling and type of diet but no significant interaction were demonstrated. Proteins and lipids content (% of dry matter) of carcasses were also measured in sampled fish. Proteins content decreased from February to November and significant differences were demonstrated between NC and HC diets. Lipids contents increased significantly throughout the experiment,but no dietary effect was demonstrated. Proteins content of viscera was also measured, and higher protein content was measured in HC neomales compared to NC neomales in August and November. Lipids content in viscera were significantly higher in HC neomales in August and November compared to NC neomales. The viscero-somatic index (VSI) decreased throughout the experiment but no difference between diets was demonstrated. The hepato-somatic index (HSI) and the protein content of liver were respectively higher and lower in HC neomales compared to NC neomales in February, June and August. Lipid content in liver was higher in HC neomales compared to NC neomales in August. Glycogen content in liver was significantly higher in HC neomales and NC neomales in February and June.

**Table 1:** Diet composition of the non-carbohydrate diet (NC) and the high-carbohydrate diet (HC).

| Ingredients (%) | NC | HC |
| --- | --- | --- |
| Fish meal<sup>1</sup> | 77.77 | 45 |
| Pre-gelatinised starch<sup>2</sup> | - | 37 |
| CPSP90<sup>3</sup> | 2 | 5 |
| Soybean meal<sup>4</sup> | 12 | - |
| Pea protein concentrated<sup>5</sup> | - | 5 |
| Fish oil<sup>6</sup> | 1.66 | 3.96 |
| Cellulosis<sup>7</sup> | 2.53 | - |
| Alginate<sup>8</sup> | 2 | 2 |
| Minerals premix<sup>9</sup> | 1 | 1 |
| Vitamins premix<sup>9</sup> | 1 | 1 |
| Astaxanthine<sup>10</sup> | 0.04 | 0.04 |
| **Proximate composition** |  |  |
| Dry matter | 93.97 | 95.84 |
| Crude proteins (%DM) | 66.63 | 41.97 |
| Gross energy (Kj/gDM) | 20.21 | 18.73 |
| Ash (%DM) | 16.81 | 10.37 |
| Carbohydrates (%DM) | 0.55 | 32.55 |
| Crude lipids (%DM) | 8.71 | 7.57 |

<sup>1</sup>Sopropêche 62126 Wimille, France, <sup>2</sup>Gelatinised corn starch roquette 62136 Wimille, France, <sup>3</sup>Sopropêche 62126 Wimille, France, <sup>4</sup>SUDOUEST Aliment 40360 Pomarez, France, <sup>5</sup>roquette 62136 Lestrem, <sup>6</sup>Sopropêche 62126 Wimille, France, <sup>7</sup>RETTENMAIER FRANCE SARL 78100 Saint Germain en Laye, France, <sup>8</sup>Louis François 77,134 Croissy Beaubourg, France, <sup>9</sup>INRAE SAAJ 78350 Jouy en Josas, France, <sup>10</sup>DSM food 59113 Seclin, France.

**Table 2:**
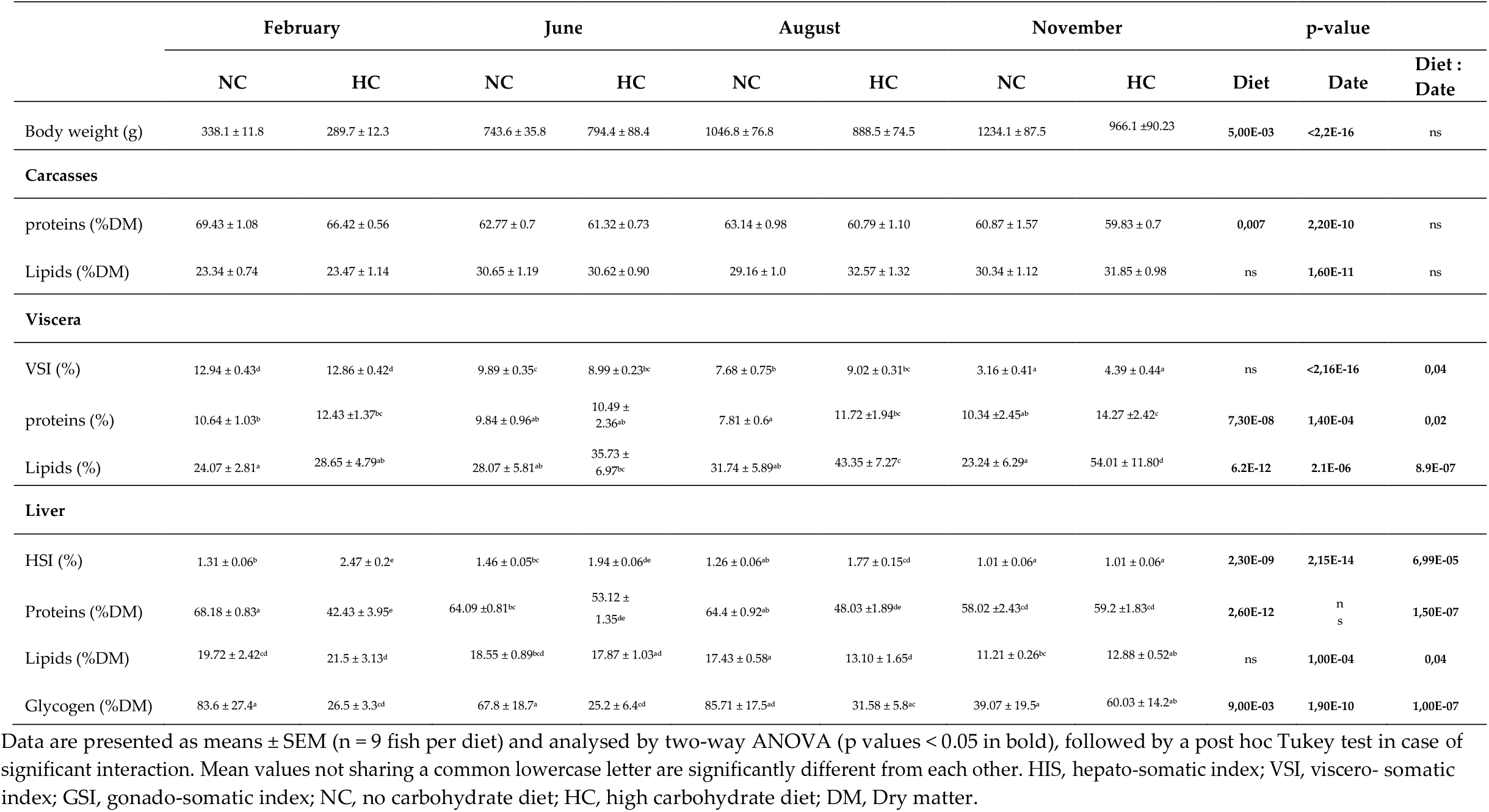
Effects of NC and HC diets on growth and biochemical composition of tissues of neomales rainbow trout broodstocks.

### 3.2 Plasma metabolites

Plasma glucose and triglycerides concentrations were analysed and compared between neomales fed with the NC diet or the HC diet during 2, 6, 8 and 11 months (figure 2). Except for August and November, plasmatic glucose concentrations were higher in neomales fed with HC diet compared to neomales fed with the NC diet. Triglycerides concentrations displayed significant differences between months of sampling, but no differences were demonstrated between diets.

**Figure 2:**
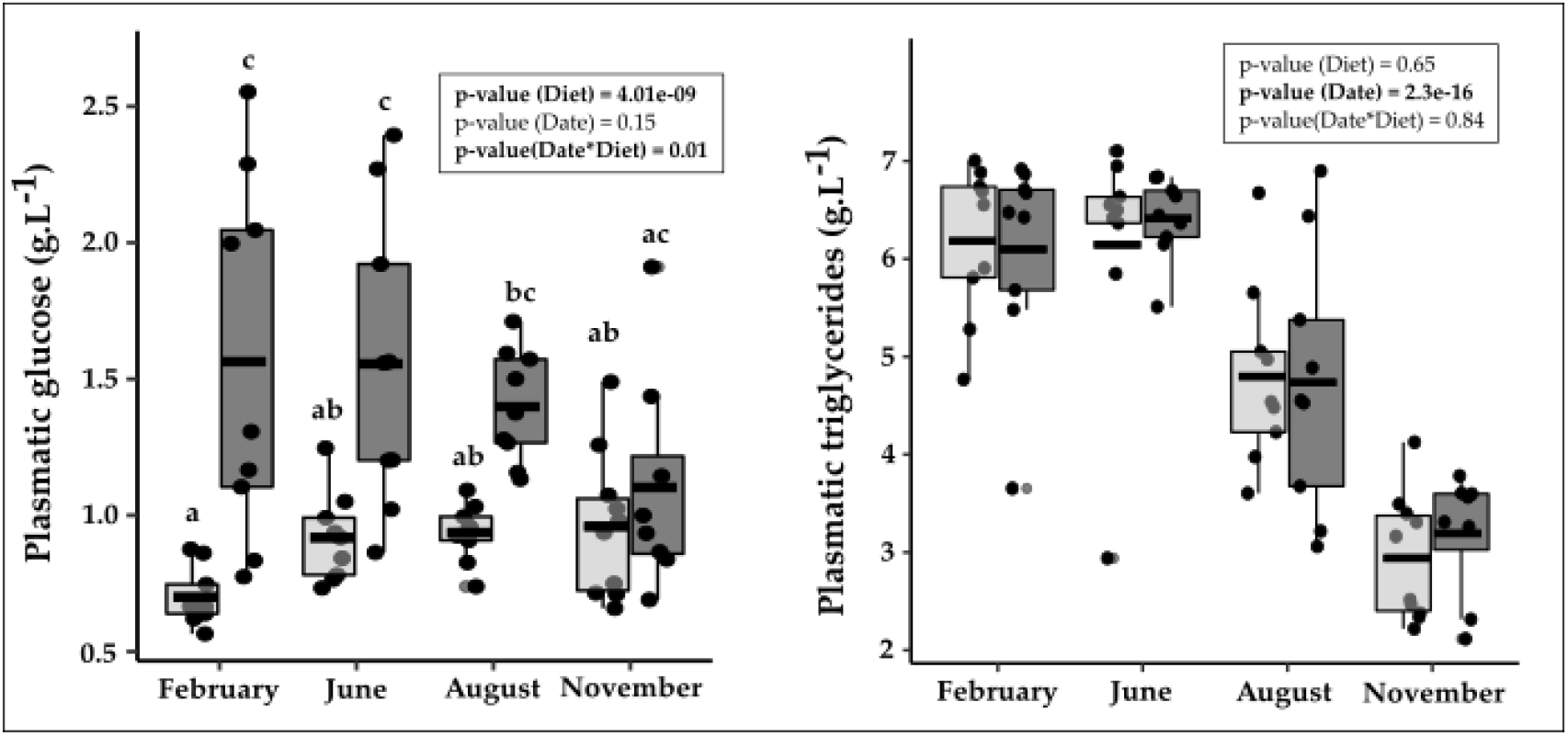
Plasma glucose and triglycerides concentrations of neomales rainbow trout fed with either the NC diet (non-carbohydrate diet, grey) or the HC diet (High carbohydrate diet, dark grey). Data (n=9 fish per diet) were analysed with a two-way ANOVA when the conditions of application were respected, followed by a post hoc Tukey’s test in case and significant differences (p < 0.05, indicated with different letters).

### 3.3 Glucose and lipid metabolism in liver

Hepatic mRNA levels of key genes in the liver involved in the glucose transport, the glycolysis, the gluconeogenesis, the glucose-6-Phosphate dehydrogenase, the *de novo* lipogenesis were investigated and compared between neomales fed with either the NC or the HC diet during 2, 6, 8 and 11 months (Table 3).

**Table 3:** Relative mRNA levels of glucose transport, glycolysis, gluconeogenesis, pentose phosphate pathway, *de novo* lipogenesis related genes in liver of neomales rainbow trout fed with a non-carbohydrate diet (NC) or a high-carbohydrate diet (HC). Data are represented as means ± SD (n = 9 neomales per diet). Genes which have not been detected are represented by “nd”. Data were analysed using a two-way ANOVA when conditions of application were respected. Significant differences of means were investigated using a post hoc Tuley’s test and are represented using different letters and bold p-values.

|  | Februray |  | June |  | August |  | November |  | p-value |  |  |
| --- | --- | --- | --- | --- | --- | --- | --- | --- | --- | --- | --- |
|  | NC | HC | NC | HC | NC | HC | NC | HC | Diet | Date | Diet : Date |
| **Glucose transport** |  |  |  |  |  |  |  |  |  |  |  |
| *glut2a* | 1.28 ± 0.2<sup>c</sup> | 0.65 ± 0.1<sup>ac</sup> | 1.18 ± 0.2<sup>ac</sup> | 0.88 ± 0.1<sup>ac</sup> | 0.57 ± 0.1<sup>a</sup> | 0.60 ± 0.1<sup>ab</sup> | 0.78 ± 0.1<sup>ac</sup> | 1.18 ± 0.2<sup>bc</sup> | ns | **2,00E-04** | **6,00E-04** |
| *glut2b* | 1.31 ± 0.1<sup>bc</sup> | 0.71 ± 0.1<sup>a</sup> | 1.27 ± 0.2<sup>bc</sup> | 0.85 ± 0.1<sup>ab</sup> | 0.88 ± 0.1<sup>ac</sup> | 0.93 ± 0.1<sup>ac</sup> | 1.10 ± 0.05<sup>ac</sup> | 1.37 ± 0.1<sup>c</sup> | **0,02** | **0,01** | **7,00E-04** |
| *glut1aa* | nd | nd | nd | nd | nd | nd | nd | nd | - | - | - |
| *glut1ab* | nd | nd | nd | nd | nd | nd | nd | nd | - | - | - |
| *glut1bb* | 1.30 ± 0.2 | 1.05 ± 0.1 | 1.18 ± 0.3 | 0.53 ± 0.1 | 1.21 ± 0.2 | 1.0 ± 0.1 | 0.92 ± 0.1 | 0.92 ± 0.2 | **4,00E-03** | **0,02** | ns |
| *glut1ba* | 0.96 ± 0.1 | 1.16 ± 0.3 | 1.13 ± 0.1 | 0.70 ± 0.1 | 0.64 ± 0.1 | 0.43 ± 0.1 | 0.59 ± 0.1 | 0.66 ± 0.1 | ns | **3,00E-03** | ns |
| **Glycolysis** |  |  |  |  |  |  |  |  |  |  |  |
| *gcka* | 0.0 ± 0.0<sup>a</sup> | 1.75 ± 0.6<sup>bc</sup> | 0.0 ± 0.0<sup>a</sup> | 1.92 ± 0.68<sup>c</sup> | 0.0 ± 0.0<sup>a</sup> | 2.07 ± 0.3<sup>c</sup> | 0.39 ± 0.2<sup>ab</sup> | 0.15 ± 0.1<sup>a</sup> | **3,70E-12** | 0.09 | **5,60E-07** |
| *gckb* | 0.003 ± 0.002<sup>a</sup> | 1.63 ± 0.6<sup>b</sup> | 0.001 ± 0.001<sup>a</sup> | 2.25 ± 0.4<sup>b</sup> | 0.0 ± 0.0<sup>a</sup> | 1.58 ± 0.2<sup>b</sup> | 0.01 ± 0.0<sup>a</sup> | 0.02 ± 0.0<sup>a</sup> | **<2,1E-16** | **2,20E-10** | **3,30E-12** |
| *pfkla* | 1.03 ± 0.2 | 1.4 ± 0.2 | 0.82 ± 0.2 | 0.68 ± 0.1 | 1.32 ± 0.1 | 1.21 ± 0.1 | 0.8 ± 0.1 | 0.8 ± 0.1 | ns | **2,00E-04** | ns |
| *pfklb* | 1.06 ± 0.2 | 1.48 ± 0.2 | 0.88 ± 0.2 | 0.74 ± 0.15 | 1.38 ± 0.1 | 1.23 ± 0.1 | 0.81 ± 0.1 | 0.86 ± 0.1 | ns | **6,00E-04** | ns |
| *pkl* | 1.48 ± 0.1 | 0.94 ± 0.1 | 1.42 ± 0.2 | 0.91 ± 0.1 | 1.16 ± 0.1 | 0.83 ± 0.04 | 0.80 ± 0.1 | 0.81 ± 0.05 | **3,00E-04** | **6,00E-03** | 0.06 |
| **Gluconeogenesis** |  |  |  |  |  |  |  |  |  |  |  |
| *g6pca* | 1.36 ± 0.1ab | 0.88 ± 0.10a | 1.54 ± 0.20ab | 0.84 ± 0.05a | 0.89 ± 0.24a | 0.98 ± 0.17a | 0.95 ± 0.08a | 2.18 ± 0.33b | ns | **2,00E-03** | **4,90E-06** |
| *g6pcb1a* | 1.23 ± 0.2 | 1.14 ± 0.3 | 0.94 ± 0.2 | 0.61 ± 0.1 | 1.67 ± 0.3 | 3.19 ± 0.7 | 3.00 ± 0.7 | 2.73 ± 0.8 | ns | **2,50E-08** | 0.06 |
| *g6pcb1b* | 1.44 ± 0.3<sup>ac</sup> | 0.57 ± 0.2<sup>ab</sup> | 1.46 ± 0.2<sup>ac</sup> | 0.3 ± 0.1<sup>a</sup> | 1.81 ± 0.53<sup>ac</sup> | 2.67 ± 0.78<sup>c</sup> | 0.93 ± 0.2<sup>ac</sup> | 2.02 ± 0.53<sup>bc</sup> | **9,00E-03** | **2,90E-05** | **5,10E-07** |
| *g6pcb2a* | nd | nd | nd | nd | nd | nd | nd | nd | - | - | - |
| *g6pcb2b* | 0.59 ± 0.1<sup>a</sup> | 2.93 ± 0.8<sup>b</sup> | 0.57 ± 0.1<sup>a</sup> | 0.59 ± 0.4<sup>a</sup> | 0.98 ± 0.4<sup>a</sup> | 0.96 ± 0.4<sup>a</sup> | 1.0 ± 0.3<sup>a</sup> | 1.49 ± 0.3<sup>ab</sup> | **0,01** | **0,01** | **0,02** |
| *fbp1a* | nd | nd | nd | nd | nd | nd | nd | nd | - | - | - |
| *fbp1b1* | 1.41 ± 0.1<sup>d</sup> | 0.70 ± 0.1<sup>ab</sup> | 1.26 ± 0.1<sup>cd</sup> | 0.91 ± 0.1<sup>bd</sup> | 0.71 ± 0.08<sup>abc</sup> | 0.62 ± 0.1<sup>ab</sup> | 0.40 ± 0.02<sup>a</sup> | 0.56 ± 0.1<sup>a</sup> | **7,00E-03** | **1,60E-08** | **5,00E-03** |
| *fbp1b2* | 1.35 ± 0.3 | 1.17 ± 0.1 | 0.88 ± 0.2 | 0.42 ± 0.1 | 1.47 ± 0.3 | 1.10 ± 0.2 | 0.68 ± 0.2 | 0.48 ± 0.2 | **9,00E-03** | **4,60E-06** | ns |
| *pck1* | 1.18 ± 0.2<sup>c</sup> | 0.12 ± 0.03<sup>ab</sup> | 0.83 ± 0.30<sup>bc</sup> | 0.02 ± 0.004<sup>a</sup> | 0.46 ± 0.18<sup>ac</sup> | 0.02 ± 0.006<sup>a</sup> | 0.15 ± 0.04<sup>ab</sup> | 0.52 ± 0.26<sup>ac</sup> | **3,30E-05** | ns | **2,00E-04** |
| *pck2a* | 1.92 ± 0.8 | 0.91 ± 0.2 | 1.49 ± 0.4 | 0.62 ± 0.2 | 1.09 ± 0.4 | 1.36 ± 0.4 | 1.87 ± 0.4 | 2.77 ± 0.6 | ns | **4,00E-03** | 0.09 |
| *pck2b* | 1.48 ± 0.1 | 1.16 ± 0.1 | 1.28 ± 0.1 | 0.95 ± 0.1 | 1.27 ± 0.1 | 1.09 ± 0.1 | 1.18 ± 0.1 | 1.03 ± 0.07 | **1,00E-03** | ns | ns |
| **Pentose phosphate Pathway** |  |  |  |  |  |  |  |  |  |  |  |
| *g6pd* | 0.57 ± 0.1<sup>ab</sup> | 2.9 ± 0.33<sup>d</sup> | 0.36 ± 0.09<sup>a</sup> | 1.3 ± 0.34<sup>bcd</sup> | 0.97 ± 0.44<sup>ac</sup> | 2.04 ± 0.21<sup>cd</sup> | 0.22 ± 0.02<sup>a</sup> | 0.29 ± 0.05<sup>a</sup> | **4,90E-08** | **2,60E-08** | **8,00E-03** |
| **De novo lipogenesis** |  |  |  |  |  |  |  |  |  |  |  |
| *srebp1* | 0.96 ± 0.2 | 2.20 ± 0.4 | 0.92 ± 0.2 | 0.82 ± 0.2 | 1.68 ± 0.4 | 1.27 ± 0.3 | 0.95 ± 0.2 | 0.47 ± 0.07 | **3,60E-07** | **9,40E-11** | ns |
| *aca α a* | 0.84 ± 0.1 | 2.28 ± 0.3 | 0.68 ± 0.2 | 1.29 ± 0.2 | 1.62 ± 0.4 | 2.45 ± 0.3 | 0.76 ± 0.2 | 1.35 ± 0.3 | **4,60E-05** | **5,00E-03** | ns |
| *aca α b* | 0.28 ± 0.1<sup>ab</sup> | 2.57 ± 0.5<sup>c</sup> | 0.46 ± 0.4<sup>ab</sup> | 0.53 ± 0.2<sup>b</sup> | 4.63 ± 1.4<sup>c</sup> | 9.78 ± 2.3<sup>c</sup> | 0.16 ± 0.04<sup>ab</sup> | 0.08 ± 0.02<sup>a</sup> | **2,00E-03** | **2,00E-15** | **3,00E-04** |
| *acly* | 0.47 ± 0.1 | 2.46 ± 0.6 | 0.42 ± 0.1 | 1.67 ± 0.6 | 0.9 ± 0.4 | 2.24 ± 0.7 | 0.11 ± 0.01 | 0.20 ± 0.05 | **3,60E-07** | **9,40E-11** | ns |
| *fasn* | 0.81 ± 0.1 | 2.35 ± 0.4 | 0.74 ± 0.2 | 1.19 ± 0.4 | 2.2 ± 0.8 | 3.07 ± 0.6 | 0.08 ± 0.02 | 0.05 ± 0.01 | **0,04** | **<2E-16** | 0.09 |

### 3.4 Glucose transport

For the transport of glucose, *glut2a* displayed a significant interaction (Date : Diet) with lower mRNA levels for neomales fed with HC or the NC diet in August compared to neomales fed with the NC diet in February (Table 3). *glut2b* displayed also a significant interaction with lower mRNA levels in HC neomales in February compared to NC neomales in February, in June and HC neomales in November. Regarding the *glut1-* related genes only *glut1bb* and *glut1ba* were detected displaying significant differences between diet and date for *glut1bb* and date for *glut1ba*.

### 3.5 Glycolysis

For the glycolysis, *gcka* and *gckb* displayed a significant interaction with higher mRNA levels of both genes in neomales fed with the HC diet compared to neomales fed with the NC diet in February, June and August. No differences were observed in fasted neomales in November. *pfkla, pfklb* and *pkl* displayed no significant interaction but significant differences were demonstrated between dates. *pkl* also displayed significant differences among groups of diet.

### 3.6 Gluconeogenesis

For the gluconeogenesis, *g6pca, g6pcb1b, g6pcb2b, fbp1b1* and *pck1* displayed a significant interaction between date and diet (Table 3). *g6pca* showed higher mRNA levels in HC neomales compared to NC neomales in November and August as well as compared to HC neomales in February, June and August. *g6pcb1b* displayed higher mRNA levels in HC neomales in August compared to HC neomales in February and June. *g6pcb2b* displayed also higher mRNA levels in HC neomales in February compared to NC neomales but no significant differences between diets were demonstrated for other date. *fbp1b1* displayed higher mRNA levels in NC neomales compared to HC neomales in February but no differences were demonstrated between NC and HC diets for other months. *pck1* displayed higher mRNA levels in NC neomales compared to HC neomales in February and June but not in August and November. *g6pcb1a* and *pck2a* displayed significant differences between date while *fbp1b2* displayed significant differences between date and diet and *pck2b* significant differences between diets.

### 3.7 The *de novo* lipogenesis and the glucose-6-phosphate dehydrogenase

The *de novo* lipogenesis and the glucose-6-phosphate dehydrogenase related genes were investigated throughout the reproductive cycle of neomales fed with a NC diet or a HC diet (Table 3). *g6pd* and *aca alpha b* displayed significant interaction between date and diet. *g6pd* displayed higher mRNA levels in HC neomales compared to NC neomales both in February and June. *aca alpha b* displayed higher mRNA levels in HC neomales compared to NC neomales but only in February. *srebp1, aca alpha a, acly* and *fasn* displayed no significant interaction but the effect of both date and diet was demonstrated.

### 3.8 Enzymatic activities in the liver

Enzymatic activities were analysed and compared between neomales fed with the NC diet and neomales fed with the HC diet during, 2, 4, 6 and 8 months (respectively February, June, August, November) (figure 3). For the glycolysis, only Gck displayed a significant interaction between date and diet with, except in November, higher hepatic activities in neomales fed with the HC diet compared to neomales fed with the NC diet. Pfk enzymatic assay demonstrated no significant differences between NC and HC neomales in February, June, August and November but significant effect of date was demonstrated. Pk displayed significant effects of date and diet. For the gluconeogenesis enzymes, no significant interaction between date and diet were demonstrated. A significant effect of date was demonstrated for G6pc, Fbp and Pck activities and except for Fbp, a significant effect of diet was also demonstrated. For the pentose phosphate pathway, G6pd displayed significant effect of date and diet but no significant interaction. For the *de novo* lipogenesis, Fasn displayed significant differences between dates (February, June, August, November).

**Figure 3:**
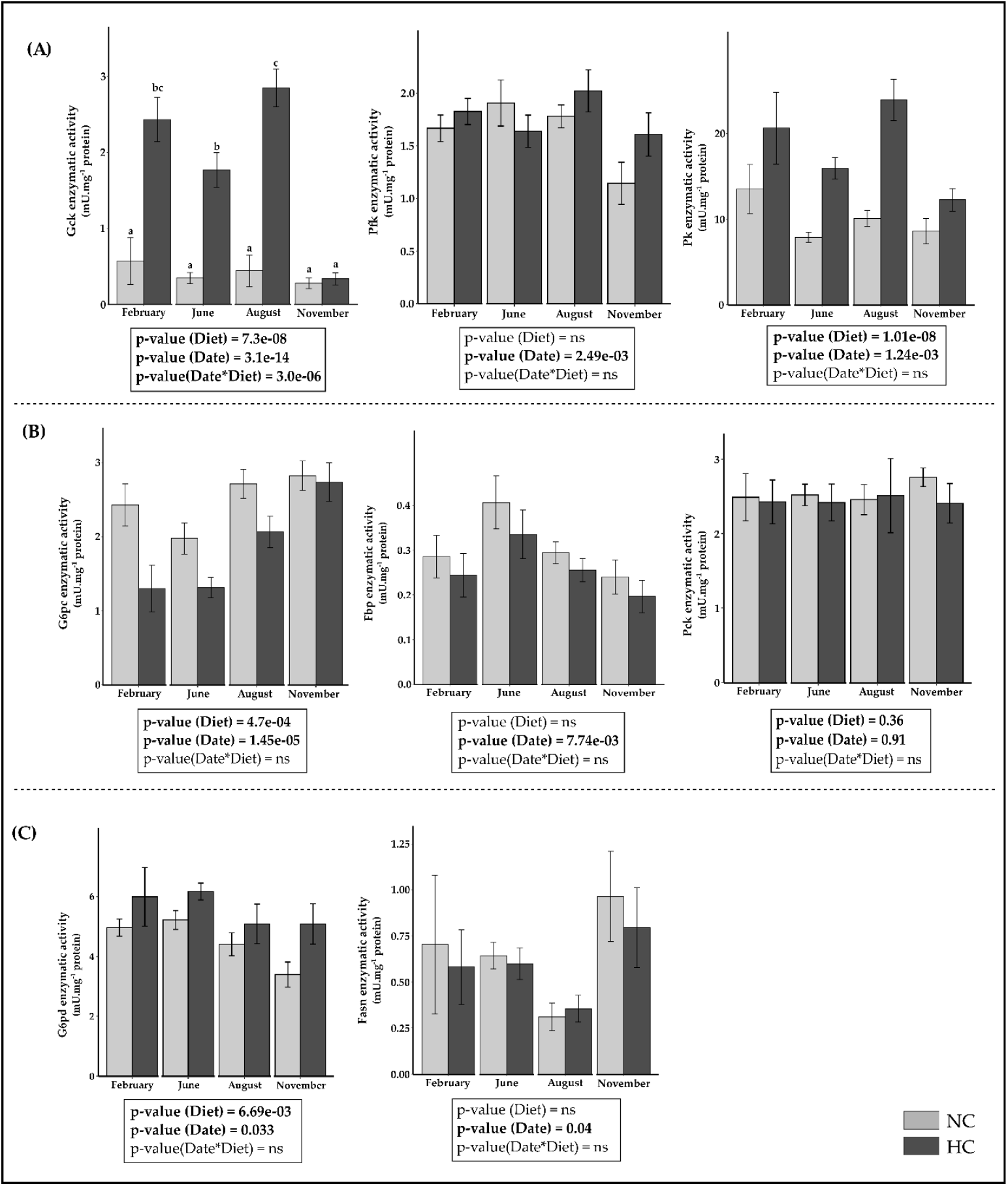
Enzymatic activities of (A) glycolysis, (B) gluconeogenesis and (C) pentose phosphate pathway and *de novo* lipogenesis in the liver of neomales rainbow trout fed with a non-carbohydrate diet (NC) or a high-carbohydrate diet (HC). Enzymatic assays are expressed as mU.mg-1 of proteins for (A) the glucokinase (Gck), the phosphofructokinase (Pfk) and the pyruvate kinase (Pk), (B) the glucose-6-phosphatase (G6pc), the fructose-1,6-bisphosphatase (Fbp) and the phosphoenolpyruvate carboxykinase (Pck) and (C) the glucose-6-phosphate dehydrogenase (G6pd) and the fatty acid synthase (Fasn). Data were analysed using a two-way ANOVA when conditions of application were respected. Significant differences of means were investigated using a post hoc Tuley’s test and are represented using different letters and bold p-values.

### 3.9 Reproductive performances

Reproductive performances were assessed and compared through survival at eyed-up stage, at hatching and percentages of malformation in the progeny of NC fed females cross-fertilised with milts of NC or HC neomales (figure 4). To mitigate the potential maternal effect, we fertilised the eggs of the same females with NC and HC neomales (respectively NN and NH). Lower survival rate at eye-stage were obtained for the group NH (85.3% against 93.5% for NN) and higher survival rate at hatching were observed in NH (94.6% against 90.2% for NN). No difference of percentage of malformation was demonstrated between progeny of NC and HC neomales.

**Figure 4:**
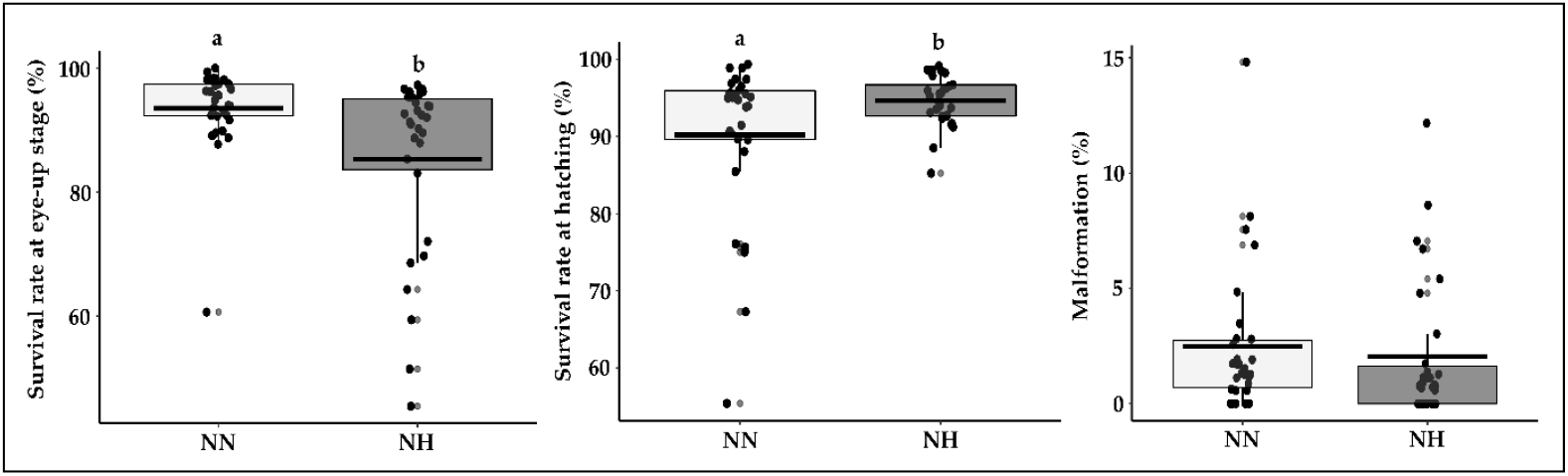
Survival rates at eye-up stage, at hatching and percentages of malformation in the progeny of females fed with a non-carbohydrate diet cross-fertilised with milts of neomales fed with the NC diet (NN – grey) or fed with the HC diet (NH – dark grey). Data (n=36 crossings per condition) were analysed by a one-way ANOVA followed by a post hoc Tukey’s test in case of significant differences (p < 0.05, indicated by different letters).

## 4 Discussion

Rainbow trout is a carnivorous fish adapted for high-protein catabolism and low-carbohydrate use. This species display a persistent postprandial hyperglycaemia when fed a diet containing more than 20% of digestible carbohydrates [1]. While this feature is well-described in juveniles less attention is given to rainbow trout broodstocks and especially neomales that are used to produce all-female population. In a previous study we were able to demonstrate that a short carbohydrate intake (two-day feeding) did not induce specific metabolic changes in the liver of females, males and neomales broodstocks [13]. Moreover, we also demonstrated that neomales displayed more pronounced modifications of glucose and lipid metabolisms in response to nutritional status or HC diet compared to both females and males broodstocks. Indeed, neomales displayed enhanced glycolysis potential compared to both females and males and differences in nutritional regulation of mRNA levels irrespective of the metabolism investigated. Finally, a long-term feeding trial on females and males rainbow trout broodstock revealed that they were able to eat and use a high carbohydrate diet without adverse effect on reproduction [26]. Thus, it appears relevant to test, for the first time, long-term effects of feeding neomales with a high-carbohydrate diet on their growth and reproductive performances as well as assessing metabolic changes in the liver and gonads.

### 4.1 Neomales can eat and grow with a high carbohydrate diet during a complete reproductive cycle

Dietary carbohydrate intake leads to the increase of expression and activity of Gck in trout [27]. Except for neomales sampled in November that were fasted for 10 days, we observed an increase of *gcka* and *gckb* mRNA levels and activities in HC neomales confirming that neomales in our study ate and metabolized dietary carbohydrates. Moreover, all HC neomales sampled were also hyperglycaemic 6h after the last meal confirming the ingestion of glucose as well as its storage in the form of glycogen in the liver with higher proportion in HC neomales compared to NC neomales. Finally, HC neomales stored glucose in the form of lipids with strong increased proportions of lipids in viscera in HC neomales compared to NC neomales in both August and November. This demonstrate that neomales are able to metabolise and store excess glucose during a long-term feeding trial with a high carbohydrate diet as well as what is observed in females and males broodstocks (Callet et al., 2020). Although HC neomales did metabolise the HC diet, the type of diet throughout the experiment did not affect FI. Nonetheless, we observed an effect of the diet on growth performances with NC neomales growing slightly better than HC neomales. This could be explained by the slight decrease of FE in HC neomales compared to NC neomales except in February and August suggesting a differential ability to eat and grow in presence of 33% of carbohydrates in the diet. In fact in our study, neomales displayed a decrease in growth performance which is observed in juveniles when fed with a HC diet and representing one feature of the so-called glucose intolerant phenotype [6,28,29]. Finally, Callet et al. (2020) [30] previously revealed that males and females broodstocks fed with a high-carbohydrate diet (32%) during respectively 4 and 8 months displayed glycaemia within a normal range suggesting that glucose homeostasis in neomales is regulated in different manner compared to both males and females in presence of dietary carbohydrates.

### 4.2 Neomales catabolise and store dietary glucose in liver depending on the reproductive cycle period

Changes in the ability to use glucose during gametogenesis were suggested to be part of the response of females and males to HC diet with an initial high energy demand in both liver and gonads [26]. This high energy requirement was represented by high glycolysis potential and increased G6pd activity in both liver and gonads accompanied by a low hepatic gluconeogenesis and followed by a drop of both hepatic Gck and G6pd enzymatic activities latter in the reproductive cycle [26]. Interestingly, we did observe a strong induction of hepatic glycolysis through enhanced *gcka* and *gckb* mRNA levels and increased levels of mRNA and activity of G6pd but not followed latter by a drop of activity of both Gck and G6pd at the end of the experiment suggesting different ability to use glucose in neomales compared to females and males during gametogenesis. Concomitant to this, and still regarding the production of energy through glycolysis, we also observed an increase in the activity of the hepatic pyruvate kinase confirming that carbohydrates are metabolised. Nonetheless, we also observed a decrease of mRNA levels of *pkl* in HC neomales that have been already described in rainbow trout juveniles and suspected to be involved in the poor glycaemia control [31]. Interestingly, the effect of diet on *pkl* mRNA levels was not demonstrated in females and males broodstocks suggesting that this feature is specific to both juveniles and neomales [26]. This could in fact justify the detection of hyperglycaemia in HC neomales compared to both females and males fed with a HC diet.

When glucose is in excess after ingestion of a high carbohydrate diet, it can be also converted into triglycerides through the *de novo* lipogenesis [6]. G6pd dictate the flux of the Pentose Phosphate Pathway that permits the production of NADPH, H+. This reducing power could be thus used by the *de novo* lipogenesis (DNL) to produce triglycerides. Our data revealed that DNL was indeed induced in presence of carbohydrates early in the reproductive cycle with an increased expression of *aca alpha b* in HC neomales even though triglycerides plasmatic levels were not affected by the type of diet. Thus, this suggest that the glucose metabolised from the HC diet could be stored into triglycerides through *de novo* lipogenesis. Interestingly, this increased expression in HC neomales compared to NC neomales was not observed later in the experiment suggesting that rather than only being regulated by the carbohydrate intake DNL could be also subject to metabolic modulation during gametogenesis in neomales as previously described in both females and males [26,32,33].

### 4.3 The hepatic glucose production is dependent on the type of sex

While glycolysis was clearly induced in HC neomales, we also demonstrated a down-regulation of the gluconeogenesis early in the reproductive cycle (February and June), except for the last step that was regulated as what is observed in juveniles. Indeed, a specific regulation of *g6pc*-duplicated genes by carbohydrates have been demonstrated in rainbow trout juveniles with increased expression of *g6pcb2*-ohnologs while *g6pca* and *g6pcb1*-ohnologs were down-regulated in presence of carbohydrates [4]. This atypical regulation compared to mammals was partially observed in our study with an increase of *g6pcb2b* mRNA levels in HC neomales but only in February suggesting that *g6pc*-related genes regulation is also modulated throughout the reproductive cycle and irrespective of the type of diet in neomales. Interestingly, both *g6pcb2a* and *g6pcb2b* were not induced in rainbow trout males broodstocks but in females after two months of feeding with a high-carbohydrate diet suggesting that even if neomales are phenotypic males they did not regulate gluconeogenesis in the same way as males early in the reproductive cycle. Surprisingly, we also observed a drastic increase of *g6pca* mRNA levels in November in fasted neomales that were previously fed with the HC diet compared to neomales that were fed with the NC diet which is again a common feature with females [26]. This highlights a higher ability to use glucose in HC neomales and in females fed with a high carbohydrate diet at the end of the reproductive cycle as well as suggesting a genotype dependent regulation rather than a phenotype dependent regulation of this pathway in rainbow trout. All this data reinforces the interest on the role of this pathway in the establishment of a glucose intolerant phenotype in rainbow trout especially through the non-inhibition of some gluconeogenic genes in presence of carbohydrates. Even though this specific regulation was described in the beginning of our experiment, we did observe metabolic changes irrespective of the diet throughout the entire reproductive cycle supporting a potential better ability to regulate gluconeogenesis during gametogenesis in neomales as well as what is observed in females [26,32,33].

### 4.4 Reproductive performance was affected by the HC diet in neomales confirming the importance of the nutrition of brood stock

Reproductive performances (survival at eye-up stage, at hatching and rate of malformation) were affected by feeding neomales with the HC diet only at eye-up stage and at hatching. Interestingly, feeding neomales with the HC diet lowered the survival rate at eye-up stage but increase the survival rate at hatching suggesting that the early development can be affected by the nutritional history of neomales. Even though the presence of carbohydrates did affect progeny’s survival these rates were relatively high and in line with previous results obtained in our facilities (from 85.3% to 93.5% at eye-up stage and for 90.2% to 94.6% at hatching). Overall, we can suggest that feeding neomales with a HC diet did not induce adverse effect on reproductive performances even though evidence point that neomales’ nutrition could play a key role in the survival of progeny.

## 5 Conclusion

Altogether, our data on the regulation of glucose and lipid metabolism by both diet and the reproductive cycle suggest that neomales display specific metabolic and physiological changes when fed with a high carbohydrate diet compared to both juveniles and females and males broodstocks. While most of the feature presented before tend to support similar responses in neomales compared to juveniles (decreased growth performance, non-inhibition of some genes of the gluconeogenesis), it appears clear that gametogenesis - as for females and males broodstocks - modulate the fate of glucose in neomales especially through regulations at the molecular level. However, it is important to note that neomales are only used for one reproductive cycle. Thus, they were two-year old and did not undergo a complete reproductive cycle before starting our study while both males and females broodstocks previously investigated by Callet et al. (2020) [30] were three-year old and already undergo a complete reproductive cycle. We can thus hypothesize that the settlement of metabolic regulation by gametogenesis may differ for the first appearance of gonads. Further studies are thus needed to work on the interactions between the first appearance of gonads and glucose metabolism. Altogether, this data still highlights the relevance to formulate specific diets in accordance with each type of broodstock displaying specificities (*i*.*e*. females, males and neomales).

## Supporting information

spllementary material

## Acknowledgements

We thank T. Callet for her indispensable help with sampling. We also thank L. for the preparation of the diets.

## References

1. Kamalam, B.S.; Medale, F.; Panserat, S. Utilisation of Dietary Carbohydrates in Farmed Fishes: New Insights on Influencing Factors, Biological Limitations and Future Strategies. Aquaculture 2017, 467, 3–27, doi:10.1016/j.aquaculture.2016.02.007.

2. Hemre, G.-I.; Mommsen, T. p.; Krogdahl, Å. Carbohydrates in Fish Nutrition: Effects on Growth, Glucose Metabolism and Hepatic Enzymes. Aquac. Nutr. 2002, 8, 175–194, doi:10.1046/j.1365-2095.2002.00200.x.

3. Prabu, E.; Felix, S.; Felix, N.; Ahilan, B.; Ruby, P. An Overview on Significance of Fish Nutrition in Aquaculture Industry. Int. J. Fish. Aquat. Stud. 2017, 5, 8.

4. Marandel, L.; Seiliez, I.; Véron, V.; Skiba-Cassy, S.; Panserat, S. New Insights into the Nutritional Regulation of Gluconeogenesis in Carnivorous Rainbow Trout (Oncorhynchus Mykiss): A Gene Duplication Trail. Physiol. Genomics 2015, 47, 253–263, doi:10.1152/physiolgenomics.00026.2015.

5. Panserat, S.; Skiba-Cassy, S.; Seiliez, I.; Lansard, M.; Plagnes-Juan, E.; Vachot, C.; Aguirre, P.; Larroquet, L.; Chavernac, G.; Medale, F.; et al. Metformin Improves Postprandial Glucose Homeostasis in Rainbow Trout Fed Dietary Carbohydrates: A Link with the Induction of Hepatic Lipogenic Capacities? Am. J. Physiol.-Regul. Integr. Comp. Physiol. 2009, 297, R707–R715, doi:10.1152/ajpregu.00120.2009.

6. Polakof, S.; Panserat, S.; Soengas, J.L.; Moon, T.W. Glucose Metabolism in Fish: A Review. J. Comp. Physiol. B 2012, 182, 1015–1045, doi:10.1007/s00360-012-0658-7.

7. Andoh, T. Amino Acids Are More Important Insulinotropins than Glucose in a Teleost Fish, Barfin Flounder (Verasper Moseri). Gen. Comp. Endocrinol. 2007, 151, 308–317, doi:10.1016/j.ygcen.2007.01.015.

8. Capilla, E.; Médale, F.; Navarro, I.; Panserat, S.; Vachot, C.; Kaushik, S.; Gutiérrez, J. Muscle Insulin Binding and Plasma Levels in Relation to Liver Glucokinase Activity, Glucose Metabolism and Dietary Carbohydrates in Rainbow Trout. Regul. Pept. 2003, 110, 123–132, doi:10.1016/S0167-0115(02)00212-4.

9. Díaz, M.; Capilla, E.; Planas, J.V. Physiological Regulation of Glucose Transporter (GLUT4) Protein Content in Brown Trout (Salmo Trutta) Skeletal Muscle. J. Exp. Biol. 2007, 210, 2346–2351, doi:10.1242/jeb.002857.

10. Nynca, J.; Kuźmiński, H.; Dietrich, G.J.; Hliwa, P.; Dobosz, S.; Liszewska, E.; Karol, H.; Ciereszko, A. Biochemical and Physiological Characteristics of Semen of Sex-Reversed Female Rainbow Trout (Oncorhynchus Mykiss, Walbaum). Theriogenology 2012, 77, 174–183, doi:10.1016/j.theriogenology.2011.07.039.

11. Nynca, J.; Kuźmiński, H.; Dietrich, G.J.; Hliwa, P.; Dobosz, S.; Liszewska, E.; Karol, H.; Ciereszko, A. Changes in Sperm Parameters of Sex-Reversed Female Rainbow Trout during Spawning Season in Relation to Sperm Parameters of Normal Males. Theriogenology 2012, 77, 1381–1389, doi:10.1016/j.theriogenology.2011.11.001.

12. Jouhari, S.A.; Kalbasi, M.R.; Veylaky, A.S.; Tala, M. Comparison Of Sperm Traits And Fertilization Performance Of Normal Male And Neomale Rainbow Trout (Oncorhynchus Mykiss). 2003, 2, 27–38.

13. Favalier, N.; Véron, V.; Marchand, M.; Surget, A.; Maunas, P.; Turonnet, N.; Panserat, S.; Marandel, L. Short-Term Effect of a Low-Protein High-Carbohydrate Diet on Mature Female and Male, and Neomale Rainbow Trout. Int. J. Mol. Sci. 2021, 22, 6149, doi:10.3390/ijms22116149.

14. Demska-Zakęś, K.; Hliwa, P.; Matyjewicz, P.; Zakęś, Z. The Effect of 17a-Methyltestosterone and 11b-Hydroxyandrostenedione on the Development of Reproductive System in Rainbow Trout (Oncorhynchus Mykiss Walbaum). Fish. Aquat. Life 1999, 7, 227–235.

15. Good, C.A.; Kramer, H.; Somogyi, M. The Determination of Glycogen. J. Biol. Chem. 1933, 100, 485–491.

16. Liu, J.; Dias, K.; Plagnes-Juan, E.; Veron, V.; Panserat, S.; Marandel, L. Long-Term Programming Effect of Embryonic Hypoxia Exposure and High-Carbohydrate Diet at First Feeding on Glucose Metabolism in Juvenile Rainbow Trout. J. Exp. Biol. 2017, 220, 3686–3694, doi:10.1242/jeb.161406.

17. Sambroni, E.; Lareyre, J.-J.; Gac, F.L. Fsh Controls Gene Expression in Fish Both Independently of and through Steroid Mediation. PLOS ONE 2013, 8, e76684, doi:10.1371/journal.pone.0076684.

18. Bellaiche, J.; Lareyre, J.-J.; Cauty, C.; Yano, A.; Allemand, I.; Le Gac, F. Spermatogonial Stem Cell Quest: Nanos2, Marker of a Subpopulation of Undifferentiated A Spermatogonia in Trout Testis1. Biol. Reprod. 2014, 90, 79, 1–14, doi:10.1095/biolreprod.113.116392.

19. Vandesompele, J.; De Preter, K.; Pattyn, F.; Poppe, B.; Van Roy, N.; De Paepe, A.; Speleman, F. Accurate Normalization of Real-Time Quantitative RT-PCR Data by Geometric Averaging of Multiple Internal Control Genes. Genome Biol. 2002, 3, research0034.1, doi:10.1186/gb-2002-3-7-research0034.

20. Panserat, S.; Médale, F.; Blin, C.; Brèque, J.; Vachot, C.; Plagnes-Juan, E.; Gomes, E.; Krishnamoorthy, R.; Kaushik, S. Hepatic Glucokinase Is Induced by Dietary Carbohydrates in Rainbow Trout, Gilthead Seabream, and Common Carp. Am. J. Physiol.-Regul. Integr. Comp. Physiol. 2000, 278, R1164–R1170, doi:10.1152/ajpregu.2000.278.5.R1164.

21. Tranulis, M.A.; Dregni, O.; Christophersen, B.; Krogdahl, Å.; Borrebaek, B. A Glucokinase-like Enzyme in the Liver of Atlantic Salmon (Salmo Salar). Comp. Biochem. Physiol. B Biochem. Mol. Biol. 1996, 114, 35–39, doi:10.1016/0305-0491(95)02119-1.

22. Metón, I.; Caseras, A.; Fernández, F.; Baanante, I.V. Molecular Cloning of Hepatic Glucose-6-Phosphatase Catalytic Subunit from Gilthead Sea Bream (Sparus Aurata): Response of Its MRNA Levels and Glucokinase Expression to Refeeding and Diet Composition. Comp. Biochem. Physiol. B Biochem. Mol. Biol. 2004, 138, 145–153, doi:10.1016/j.cbpc.2004.03.004.

23. Chakrabarty, K.; Leveille, G.A. Acetyl CoA Carboxylase and Fatty Acid Synthetase Activities in Liver and Adipose Tissue of Meal-Fed Rats. Proc. Soc. Exp. Biol. Med. 1969, 131, 1051–1054, doi:10.3181/00379727-131-34038.

24. Bautista José M.; Garrido-Pertierra, A.; Soler, G. Glucose-6-Phosphate Dehydrogenase from Dicentrarchus Labrax Liver: Kinetic Mechanism and Kinetics of NADPH Inhibition. Biochim. Biophys. Acta BBA - Gen. Subj. 1988, 967, 354–363, doi:10.1016/0304-4165(88)90098-0.

25. Alegre, M.; Ciudad, C.J.; Fillat, C.; Guinovart, J.J. Determination of Glucose-6-Phosphatase Activity Using the Glucose Dehydrogenase-Coupled Reaction. Anal. Biochem. 1988, 173, 185–189, doi:10.1016/0003-2697(88)90176-5.

26. Callet, T.; Hu, H.; Larroquet, L.; Surget, A.; Liu, J.; Plagnes-Juan, E.; Maunas, P.; Turonnet, N.; Mennigen, J.A.; Bobe, J.; et al. Exploring the Impact of a Low-Protein High-Carbohydrate Diet in Mature Broodstock of a Glucose-Intolerant Teleost, the Rainbow Trout. Front. Physiol. 2020, 11.

27. Panserat, S.; Médale, F.; Blin, C.; Brèque, J.; Vachot, C.; Plagnes-Juan, E.; Gomes, E.; Krishnamoorthy, R.; Kaushik, S. Hepatic Glucokinase Is Induced by Dietary Carbohydrates in Rainbow Trout, Gilthead Seabream, and Common Carp. Am. J. Physiol.-Regul. Integr. Comp. Physiol. 2000, 278, R1164–R1170, doi:10.1152/ajpregu.2000.278.5.R1164.

28. Panserat, S.; Capilla, E.; Gutierrez, J.; Frappart, P.O.; Vachot, C.; Aguirre, P.; Breque, J.; Kaushik, S. Glucokinase Is Highly Induced and Glucose-6-Phosphatase Poorly Repressed in Liver of Rainbow Trout ž Oncorhynchus Mykiss/ by a Single Meal with Glucose. 2001, 9.

29. Panserat, S.; Médale, F.; Brèque, J.; Plagnes-Juan, E.; Kaushik, S. Lack of Significant Long-Term Effect of Dietary Carbohydrates on Hepatic Glucose-6-Phosphatase Expression in Rainbow Trout (Oncorhynchus Mykiss)11The Genbank Accession Number for the Rainbow Trout G6Pase Sequence Is AF120150. J. Nutr. Biochem. 2000, 11, 22–29, doi:10.1016/S0955-2863(99)00067-4.

30. Callet, T.; Hu, H.; Larroquet, L.; Surget, A.; Liu, J.; Plagnes-Juan, E.; Maunas, P.; Turonnet, N.; Mennigen, J.A.; Bobe, J.; et al. Exploring the Impact of a Low-Protein High-Carbohydrate Diet in Mature Broodstock of a Glucose-Intolerant Teleost, the Rainbow Trout. Front. Physiol. 2020, 11, 303, doi:10.3389/fphys.2020.00303.

31. Song, X.; Marandel, L.; Dupont-Nivet, M.; Quillet, E.; Geurden, I.; Panserat, S. Hepatic Glucose Metabolic Responses to Digestible Dietary Carbohydrates in Two Isogenic Lines of Rainbow Trout. Biol. Open 2018, 7, doi:10.1242/bio.032896.

32. Barciela, P.; Soengas, J.L.; Rey, P.; Aldegunde, M.; Rozas, G. Carbohydrate Metabolism in Several Tissues of Rainbow Trout, Oncorhynchus Mykiss, Is Modified during Ovarian Recrudescence. Comp. Biochem. Physiol. Part B Comp. Biochem. 1993, 106, 943–948, doi:10.1016/0305-0491(93)90055-A.

33. Soengas, J.L.; Sanmartín, B.; Barciela, P.; Aldegunde, M.; Rozas, G. Changes in Carbohydrate Metabolism in Domesticated Rainbow Trout (Oncorhynchus Mykiss) Related to Spermatogenesis. Comp. Biochem. Physiol. Part B Comp. Biochem. 1993, 105, 665–671, doi:10.1016/0305-0491(93)90103-C.

