## Supplementary material for "Rainbow trout neomales broodstocks are able to eat and use a high carbohydrate diet during a complete reproductive cycle": spllementary material

### Supplementary data

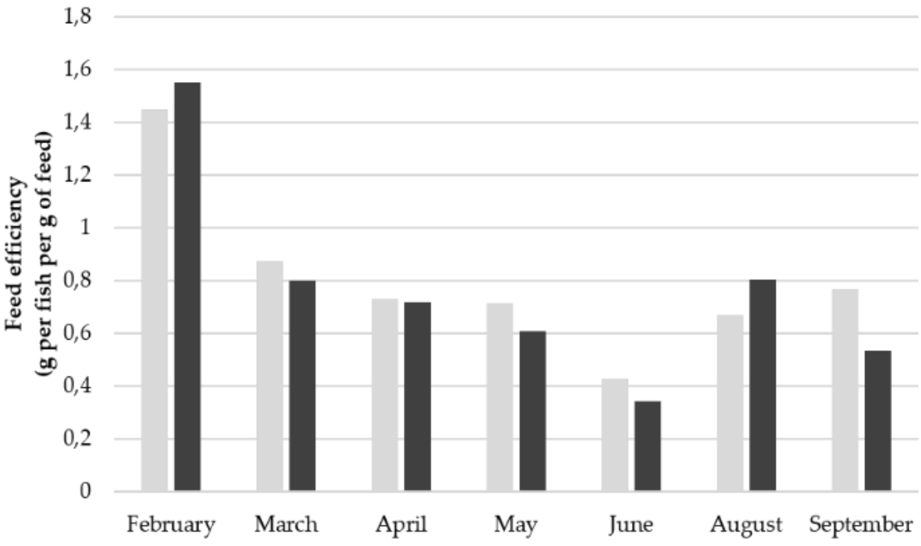


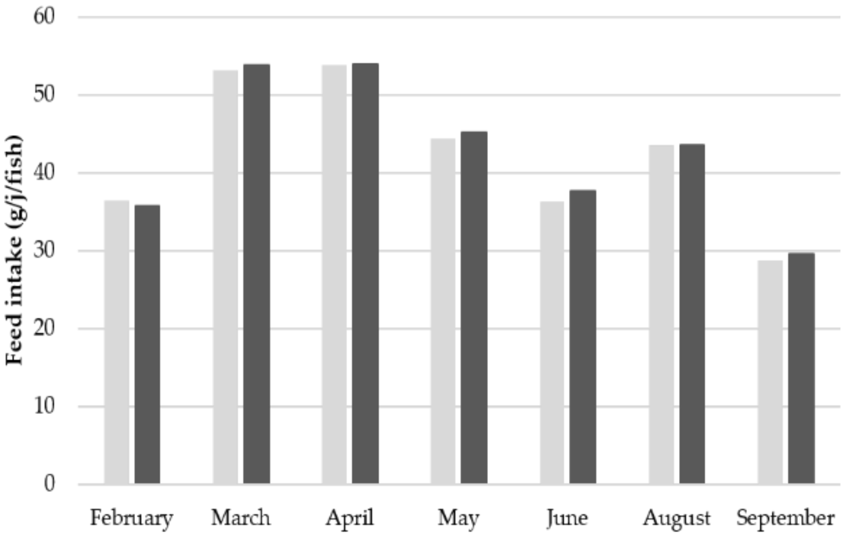


**Figure 1** : Feed efficiency and feed intake of neomales fed with a non-carbohydrate diet (NC – grey) or a high carbohydrate diet (HC – dark grey).

**Table 1 :** Effects of NC and HC diets on growth and biochemical composition in testis of neomales rainbow trout.

| | February | | June | | August | | November | | p-value | | |
|---|---|---|---|---|---|---|---|---|---|---|---|
| | NC (n = 0) | HC (n = 0) | NC (n = 7) | HC (n = 5) | NC (n = 5) | HC (n = 5) | NC (n = 5) | HC (n = 5) | Diet | Date | Diet : Date |
| GSI (%) | - | - | 0.49 ± 0.14 | 0.32 ± 0.13 | 4.55 ± 0.39 | 3.57 ± 0.83 | 3.04 ± 0.18 | 3.59 ± 0.29 | ns | **<0.05** | ns |
| Proteins (%DM) | - | - | 82.37 ± 1.66 | 80.27 ± 0.71 | 89.68 ± 2.96 | 90.38 ± 2.4 | 94.44 ± 1.4 | 91.05 ± 1.73 | ns | **<0.05** | ns |
| Lipids (%DM) | - | - | 15.47 ± 4.31 | 12.56 ± 2.30 | 10.45 ± 0.38 | 11.01 ± 0.46 | 9.53 ± 0.18 | 10.36 ± 1.21 | ns | ns | ns |

**Table 2 :** Relative mRNA levels of glucose transport, glycolysis, gluconeogenesis, steroidogenesis and spermatogenic maturation related genes in testis of neomales rainbow trout fed with a non-carbohydrate diet (NC) or a high-carbohydrate diet (HC). Data are represented as means ± SD. Genes which have not been detected are represented by "nd". Data were analysed using a two-way ANOVA when conditions of application were respected. Significant differences of means were investigated using a post hoc Tuley's test and are represented using different letters and bold p-values.

| | June | | August | | p-value | | |
|---|---|---|---|---|---|---|---|
| | NC (n = 7) | HC (n = 5) | NC (n = 5) | HC (n = 5) | Diet | Date | Diet : Date |
| **Glucose transport** | | | | | | | |
| *glut4a* | 0.09 ± 0.04 | 0.17 ± 0.05 | 0.94 ± 0.26 | 0.56 ± 0.25 | ns | **< 0.05** | ns |
| *glut4b* | nd | nd | nd | nd | - | - | - |
| *glut1aa* | nd | nd | nd | nd | - | - | - |
| *glut1ab* | nd | nd | nd | nd | - | - | - |
| *glut1bb* | 0.75 ± 0.10 | 1.04 ± 0.13 | 0.63 ± 0.05 | 0.88 ± 0.1 | **< 0.05** | ns | ns |
| *glut1ba* | 1.06 ± 0.11 | 1.23 ± 0.14 | 0.76 ± 0.23 | 1.18 ± 0.13 | 0.06 | ns | ns |
| **Glycolysis** | | | | | | | |
| *gcka* | nd | nd | nd | nd | - | - | - |
| *gckb* | nd | nd | nd | nd | - | - | - |
| *pklr* | 0.66 ± 0.22 | 1.34 ± 0.53 | 0.28 ± 0.05 | 0.30 ± 0.14 | ns | **< 0.05** | ns |
| **Gluconeogenesis** | | | | | | | |
| *g6pca* | 0.20 ± 0.1 | 0.28 ± 0.11 | 0.25 ± 0.25 | 0.66 ± 0.36 | ns | ns | ns |
| *g6pcb1a* | 0.36 ± 0.12 | 0.46 ± 0.14 | 0.57 ± 0.05 | 0.63 ± 0.16 | ns | ns | ns |
| *g6pcb1b* | 0.34 ± 0.04 | 0.21 ± 0.06 | 0.93 ± 0.31 | 0.58 ± 0.32 | ns | **< 0.05** | ns |
| *g6pcb2a* | nd | nd | nd | nd | - | - | - |
| *g6pcb2b* | nd | nd | nd | nd | - | - | - |
| *fbp1a* | 0.82 ± 0.14 | 1.17 ± 0.45 | 0.45 ± 0.18 | 0.56 ± 0.11 | ns | **< 0.05** | ns |
| *fbp1b1* | 0.48 ± 0.15 | 1.02 ± 0.42 | 0.10 ± 0.02 | 0.42 ± 0.24 | 0.06 | **< 0.05** | ns |
| *fbp1b2* | 0.74 ± 0.39 | 1.39 ± 1.01 | 0.10 ± 0.04 | 0.19 ± 0.07 | ns | ns | ns |
| *pck1* | nd | nd | nd | nd | - | - | - |
| *pck2a* | 0.60 ± 0.15 | 0.47 ± 0.06 | 0.71 ± 0.44 | 0.58 ± 0.28 | ns | ns | ns |
| *pck2b* | 0.43 ± 0.12 | 0.64 ± 0.07 | 0.42 ± 0.3 | 0.89 ± 0.14 | **< 0.05** | ns | ns |
| **Steroidogenesis** | | | | | | | |
| *fshr* | 0.56 ± 0.12 | 0.76 ± 0.13 | 1.19 ± 0.62 | 0.94 ± 0.25 | ns | ns | ns |
| *lhcgr* | 0.18 ± 0.09 | 0.50 ± 0.20 | 1.26 ± 1.26 | 0.52 ± 0.26 | ns | ns | ns |
| *star* | 0.32 ± 0.06 | 0.69 ± 0.24 | 0.97 ± 0.62 | 0.63 ± 0.15 | ns | ns | ns |
| **Spermatogenic maturation** | | | | | | | |
| *dazl* | 1.24 ± 0.19 | 1.29 ± 0.30 | 1.42 ± 0.56 | 1.47 ± 0.10 | ns | ns | ns |
| *nanos* | 0.75 ± 0.25 | 1.28 ± 0.46 | 0.28 ± 0.09 | 0.59 ± 0.26 | ns | ns | ns |
| *plzfb* | 0.36 ± 0.19 | 0.76 ± 0.28 | 0.70 ± 0.70 | 0.11 ± 0.11 | ns | ns | ns |

##### 1.1.1 Steroidogenesis and spermatogenic maturation in the testis

Spermatogenic maturation and steroidogenesis related genes were monitored throughout the experiment (Table 2 and 3). Only the sampling in June and August were compared as testis were not present in neomales in February and was largely composed of milt in November causing technicalissues in qPCR normalisation. Our results show that both steroidogenesis and spermatogenic maturation markers were not affected by the type of diet or the date. Regarding the glucose metabolism, only *pck2b* mRNA levels were affected by the diet while *glut4a*, *pkl*, *g6pcb1b*, *fbp1a* and *fbp1b1* displayed a significant effect of date.

**Table 3** *:* **:** Relative mRNA levels of glucose transport, glycolysis, gluconeogenesis, steroidogenesis and spermatogenic maturation related genes in testis of neomales rainbow trout sampled during the reproductive event in November fasted after being fed with a non-carbohydrate diet (NC) or a high-carbohydrate diet (HC). Data are represented as means ± SD.

| | November | | p-value |
|---|---|---|---|
| | NC (n = 4) | HC (n = 8) | Diet |
| **Glucose transport** | | | |
| *glut4a* | 3.2 ± 1.8 | 2.8 ± 0.8 | ns |
| *glut4b* | nd | nd | - |
| *glut1aa* | nd | nd | - |
| *glut1ab* | nd | nd | - |
| *glut1bb* | 3.2 ± 1.3 | 1.5 ± 0.29 | ns |
| *glut1ba* | 0.8 ± 0.1 | 1.27 ± 0.15 | **0.02** |
| **Glycolysis** | | | |
| *gcka* | nd | nd | - |
| *gckb* | nd | nd | - |
| *pklr* | 1.18 ± 0.32 | 2.1 ± 0.5 | ns |
| **Gluconeogenesis** | | | |
| *g6pca* | 4.84 ± 2.53 | 2.92 ± 0.92 | ns |
| *g6pcb1a* | 1.76 ± 1.0 | 2.11 ± 0.6 | ns |
| *g6pcb1b* | 0.34 ± 0.04 | 0.21 ± 0.06 | ns |
| *g6pcb2a* | nd | nd | - |
| *g6pcb2b* | nd | nd | - |
| *fbp1a* | 5.6 ± 2.3 | 0.9 ± 0.15 | **0.03** |
| *fbp1b1* | 3.4 ± 1.4 | 0.75 ± 0.17 | **0.05** |
| *fbp1b2* | 1.5 ± 0.7 | 0.75 ± 0.23 | ns |
| *pck1* | nd | nd | - |
| *pck2a* | nd | nd | - |
| *pck2b* | nd | nd | - |
| **Steroidogenesis** | | | |
| *fshr* | 8.4 ± 0.94 | 7.6 ± 1.9 | ns |
| *lhcgr* | 11.0 ± 1.1 | 9.0 ± 2.4 | ns |
| *star* | 6.4 ± 1.3 | 4.1 ± 1.05 | ns |
| **Spermatogenic maturation** | | | |
| *dazl* | 0.2 ± 0.03 | 0.3 ± 0.08 | ns |
| *nanos* | 3.9 ± 1.9 | 2.1 ± 0.5 | ns |
| *plzfb* | 3.0 ± 0.8 | 4.1 ± 1.2 | ns |
